# FANCM-family branchpoint translocases remove co-transcriptional R-loops

**DOI:** 10.1101/248161

**Authors:** Charlotte Hodson, Julienne J O’Rourke, Sylvie van Twest, Vincent J Murphy, Elyse Dunn, Andrew J Deans

## Abstract

Co-transcriptional R-loops arise from physiological or aberrant stalling of RNA polymerase, leading to formation of stable DNA:RNA hybrids. Unresolved R-loops can promote genome instability. Here, we show that the Fanconi anemia- and breast cancer-associated FANCM protein can directly unwind DNA-RNA hybrids from co-transcriptional R-loops *in vitro*. FANCM processively unwinds both short and long R-loops, irrespective of sequence, topology or coating by replication protein A. R-loops can also be unwound in the same assay by the yeast and bacterial orthologs of FANCM, Mph1 and RecG, indicating an evolutionary conserved function. Consistent with this biochemical activity of FANCM, we show that FANCM deficient cells are sensitive to drugs that stabilize R-loop formation. **Our work reveals a mechanistic basis for R-loop metabolism that is critical for genome stability.**

## Introduction

R-loops form when RNA anneals within duplex DNA, and displaces a corresponding single-strand DNA (ssDNA) patch. R-loops can arise directly, from transcription of difficult to transcribe regions, or by enzyme driven integration of RNA, such as R-loops created by the Cas9 protein during CRISPR (Ginno *et al*, 2012, Szczelkun *et al*, 2014). Persistent R-loops can be a threat to genome stability because the displaced ssDNA within an R-loop is prone to cleavage by nucleases (creating DNA breaks)(Arudchandran *et al*, 2004), recombination with distant DNA sequences (creating chromosome rearrangements)(Huertas & Aguilera, 2003) and atypical modification by ssDNA viral defence proteins such as APOBEC enzymes (creating base substitutions)(Sollier & Cimprich, 2015).

Several studies have shown DNA replication-dependent genome instability is partially prevented when the rate of transcription is reduced, indicating that R-loops cause DNA damage predominantly during S phase (Hamperl *et al*, 2017, Helmrich *et al*, 2011, Schwab *et al*, 2015). In particular, the Fanconi anemia (FA) DNA repair pathway is activated by R-loop accumulation, culminating in formation of monoubiquitinated FANCD2 at R-loop rich regions (Garcia-Rubio *et al*, 2015, Madireddy et al, 2016, Schwab *et al*, 2015). As FANCD2 monoubiquitination is normally activated by direct barriers to the replication machinery (such as DNA double strand breaks or interstrand crosslinks) (Deans & West, 2011), R-loops most likely also activate the FA pathway by blocking DNA replication. In support of this hypothesis, accumulation of monoubiquitinated FANCD2 during normal replication (but in the absence of exogenous DNA damage) is suppressed by over-expression of RNAseH1 (Madireddy *et al*, 2016, Schwab *et al*, 2015). This nuclease specifically removes DNA:RNA hybrids in the nucleus (Nakamura *et al*, 1991).

FANCM is a component of the FA pathway that is essential for activation of FANCD2 ubiquitination (Coulthard *et al*, 2013), but also has additional FA pathway-independent functions. These include direct remodeling of DNA replication fork structures, recruitment of DNA repair complexes, activation of the ATR checkpoint pathway and suppression of meiotic crossovers (Collis *et al*, 2008, Crismani *et al*, 2012, Deans & West, 2009, Gari *et al*, 2008a). We recently reported elevated formation of R-loops in FANCM-defective cells (Schwab *et al*, 2015). This was not because of increased transcription, but because of failure to properly remove R-loops (Schwab *et al*, 2015). Similarly, R-loops accumulate at telomeres after FANCM depletion, particularly in cells that utilize the ALT pathway of telomere maintenance(Pan *et al*, 2017). As such, FANCM may also prevent accumulation of DNA:RNA hybrids by the TERRA long non-coding RNA, which is essential for maintenance of ALT (Arora *et al*, 2014).

FANCM is a functional ortholog of the yeast Mph1 and Fml1 branchpoint translocase proteins (Whitby, 2010). Like these enzymes, FANCM contains an N-terminal SF2 helicase domain whose ATPase activity is activated only by branched DNA molecules (such as those found at stalled DNA replication or transcription bubbles)(Coulthard *et al*, 2013), and can translocate replication forks and Holliday junction DNA structures. Here we show for the first time that FANCM, and its yeast homolog Mph1, are also efficient in R-loop processing. Like for the bacterial RecG protein (Hong *et al*, 1995, Vincent *et al*, 1996), this function depends upon ATP hydrolysis and involves branch migration, to directly remove RNA trapped in co-transcriptionally formed R-loops.

FANCM can unwind R-loops of different size and sequence, including a highly stable R-loop formed by transcription across a telomeric repeat. Importantly, we also show that FANCM deficient cells, but not those lacking another FA group member, are highly sensitive to small molecule drugs that promote R-loop formation.

## Results

### *In vitro* unwinding of co-transcriptional R-loops by the FANCM-FAAP24 complex

We previously showed that FANCM can unwind DNA:RNA hybrids within a homologous duplex (Schwab *et al*, 2015), a property it shares with other human enzymes such as BLM and replicative helicases (Chang *et al*, 2017, Shin & Kelman, 2006). Because this type of structure may not accurately represent the formation of a co-transcriptional R-loop, we set about establishing a more sophisticated substrate that contains the high GC skew, closed triplex structure and longer length of native R-loops (Ginno *et al*, 2013). To do this we used a pUC19 plasmid containing the mouse immunoglobulin class switch recombination sequence (sµ region) in between a T7 promotor and terminator (Supplemental Figure 1a). Using T7 polymerase and ^32^P-UTP we generated co-transcriptional R-loops using techniques previously described by Roy et al (Roy *et al*, 2008), where the formation of R-loops can be measured by both a change in plasmid mobility, and retention of the radiolabeled nascent RNA (Figure 1a). Treatment with RnaseH but not Rnase A led to loss of the R-loop structure on the gel, and loss of signal on autoradiographs, confirming the presence of RNA-DNA hybrids within the plasmids (Figure 1b lane4). RNase H degraded the RNA molecule down to nucleotide sized fragments, however addition of FANCM (in heterodimer with its stabilization partner FAAP24) led to release of the RNA transcript without degradation (Figure 1c). This process was ATP dependent: the R-loop remained intact when wild-type FANCM:FAAP24 was added in the absence of ATP (Figure 1c, lane 3) or when the ATPase dead FANCM mutant complex FANCM^K117R^-FAAP24 was used (Figure 1c). We also tested whether other members of the FA pathway could act on R-loops, given that *in vivo* studies suggest the entire FA pathway (Supplemental Figure 2a) may be required for R-loop regulation (Garcia-Rubio *et al*, 2015, Schwab *et al*, 2015). No component of the FA core complex of proteins, or the FANCI:FANCD2 heterodimer had any direct effect on co-transcriptional R-loops in our assay. Under these experimental conditions only the FANCM-FAAP24 complex can unwind an R-loop structure. (Supplemental Figure 2b).

**Figure 1:**
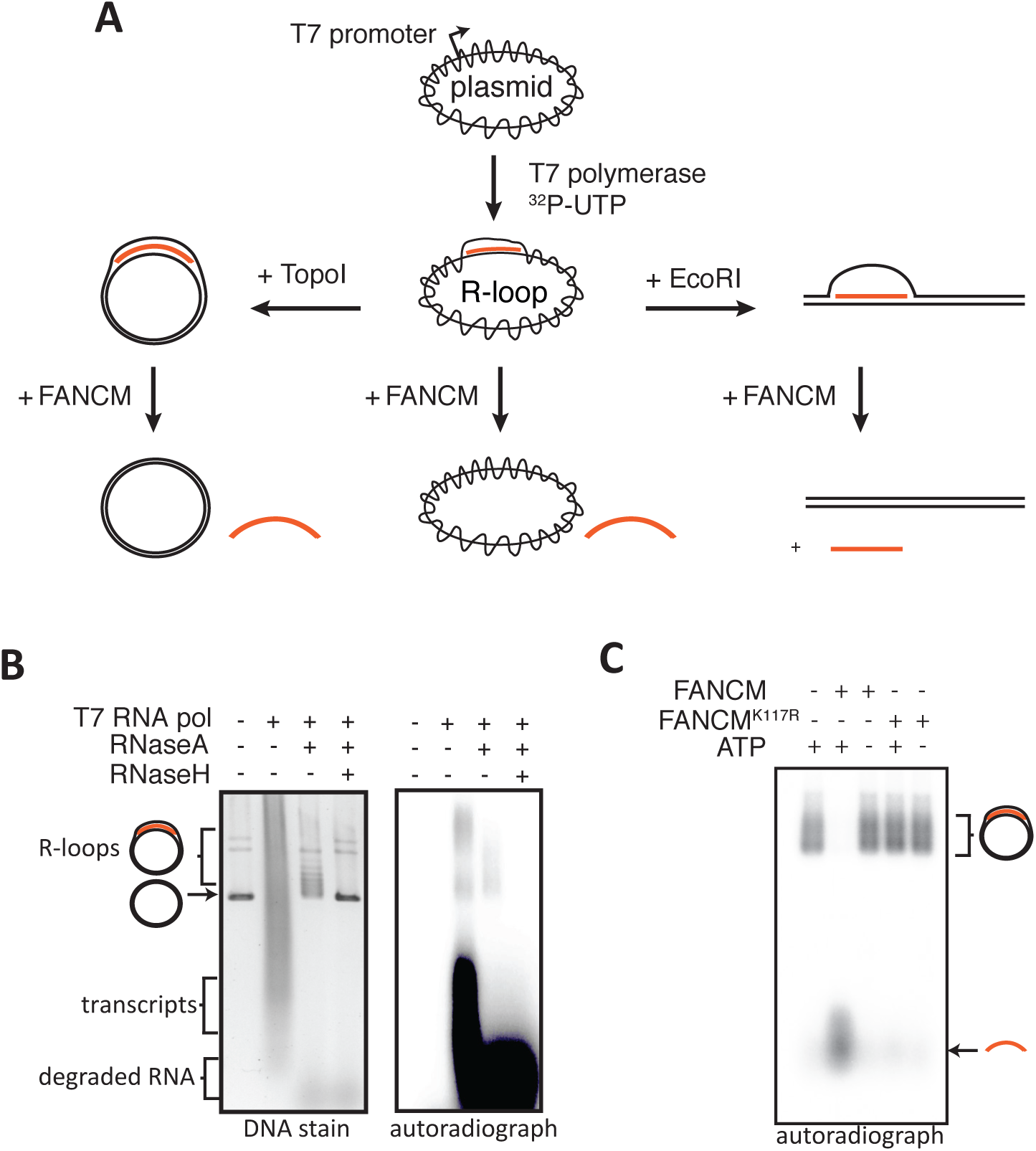
Generation of co-transcriptional R-loops in vitro and their processing by FANCM-FAAP24. A) Schematic of method used to generate and unwind R-loops of different topological states. DNA is coloured black and RNA orange. B) Plasmid based R-loops observed by gel electrophoresis. Sybr gold stain of plasmid DNA molecules reveals the topological changes to the plasmid DNA upon R-loop formation. Right panel is an autoradiograph identifying the 32P-UTP incorporation into R-loop, transcripts or post-RNAse treatment. C) Autoradiograph showing FANCM-FAAP24 (1nM) unwinding purified R-loops (1nM) in an ATP dependent manner. FANCMK117R is a translocase activity deficient mutant.

**Figure 2:**
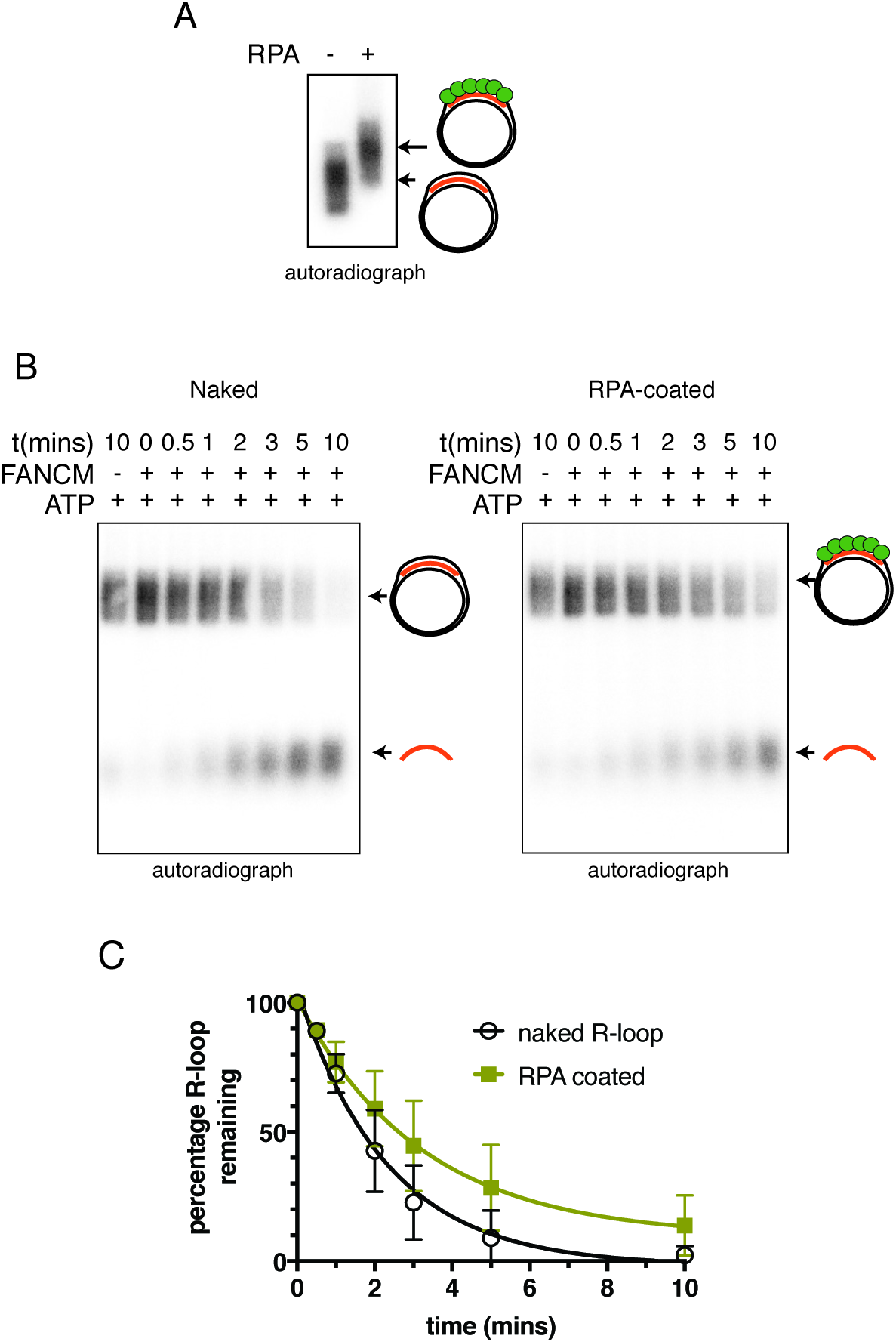
FANCM-FAAP24 acts on protein coated R-loops. A) Autoradiograph of an EMSA showing RPA bound to plasmid based R-loops. B) Representative Autoradiographs of time course assays of FANCM (0.25nM) activity on naked R-loops (1nM, left panel) versus RPA coated R-loops (1nM right panel) C) Quantification of unwinding activity (± stderr) from n>3 experiments.

### FANCM-FAAP24 unwinding of co-transcriptional R-loops is processive and not blocked by RPA binding

*In vivo*, ssDNA displaced within an R-loop is most likely bound by ssDNA binding proteins such as Replication Protein A (RPA), to protect it from attack by DNA modifying enzymes (Nguyen *et al*, 2017). To determine whether FANCM-FAAP24 could still unwind R-loops in which the displaced strand is coated in RPA filament, we incubated plasmid R-loops with molar excess of RPA for 15 minutes prior to adding FANCM-FAAP24. As expected, a uniform shift in electrophoretic mobility was observed when the displaced DNA strand in the R-loop became bound by RPA (Figure 2a). After addition of FANCM-FAAP24, 70% of the RPA bound R-loops were unwound within 5 minutes, whereas 92% of uncoated R-loops were unwound. By 10 minutes, almost all of the uncoated R-loops and approximately 85% of RPA bound R-loops were unbound (Figure 2b–c). This observation was not due to addition of extra protein to the reaction because high concentrations of Bovine Serum Albumin (BSA) did not cause an effect (Supplemental Figure 3). Together, these data suggest FANCM-FAAP24 acts in a processive manner on both naked and RPA-coated R-loops, and that RPA has a slight inhibitory effect on the unwinding ability of FANCM.

**Figure 3:**
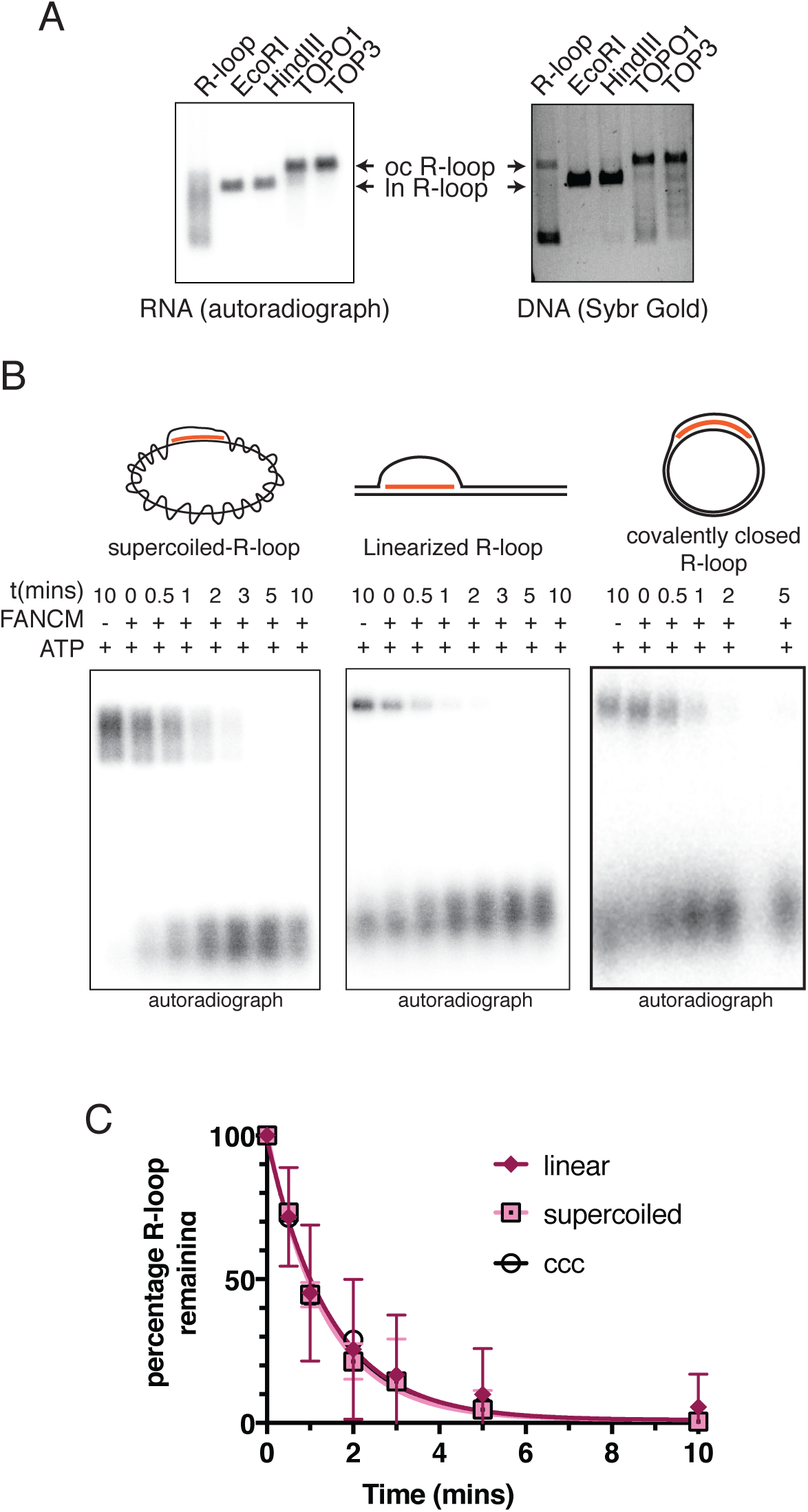
FANCM unwinds R-loops with different DNA topologies. A) Characterization of topological isoforms of plasmid based R-loops after generation of linear form by restriction enzyme digestion by EcoRI or HindIII, or covalently closed circle form (ccc) by topoisomerases 1 or III treatment. Left panel is an autoradiograph depicting the RNA molecules. The right panel is sybr gold stain of the DNA molecules. B) Autoradiographs of time course assays of plasmid (left panel) versus linear (middle panel) based R-loops. The graph is the average (n=3) % of R-loop unwound (y-axis) of each time point (x-axis) with standard error bars. Final concentrations of R-loops and FANCM for these assays was 1nM and 0.25nM respectively.

### FANCM-FAAP24 can unwind R-loops of different topology

Topological changes in DNA occur during the formation of R-loops, because the bound RNA creates underwound or overwound (supercoiled) regions. Topological effects may both promote R-loop formation during transcription (El Hage *et al*, 2010, Powell *et al*, 2013) or prevent R-loop removal, including by FANCM or other enzymes. To test this, we first assessed whether our purified plasmid R-loops were stable upon induction of topological change. As linearization removes all covalent topology from DNA, we treated our plasmid R-loops with restriction enzymes *Eco*RI and *Hind*III. We found that linearization had no effect on R-loop stability (Figure 3a). Second, R-loops were also stable in a un-super-coiled covalently closed circular (ccc) plasmid generated by *E.coli* Topo1 or human Topoisomerase IIIα:RMI1:RMI2 complex, with which FANCM associates in cells (Deans & West, 2009) (Figure 3a). Together, these data show that once formed, transcriptional R-loop structures are stable regardless of changes in DNA topology. These observations and others (Wilson-Sali & Hsieh, 2002) support the idea that type I topoisomerase role in transcription is in prevention of R-loop formation, but not the direct removal of RNA trapped within an R-loop.

We next tested the ability of FANCM-FAAP24 to unwind the topologically distinct R-loop forms. In time course assays, FANCM showed no preference in activity towards either supercoiled, linear or ccc R-loops (Figure 3b–c) unwinding all at essentially equal rates. This suggests that DNA topology does not affect R-loop unwinding activity of FANCM against a native R-loop structure.

### FANCM-FAAP24 R-loop unwinding activity is sequence and length independent

R-loops accumulate in different regions of the genome including highly transcribed genes, GC-skewed promoters and telomeric repeats (Arora *et al*, 2014, Ginno *et al*, 2012, Powell *et al*, 2013). To test whether R-loop processing by FANCM showed any sequence preference, we tested several sequences that were previously demonstrated to be strongly R-loop prone (Ginno *et al*, 2012). These sequences include the human *APOE* or *SNRPN* and mouse *Airn* genomic loci, which we cloned into our *in vitro* R-loops test plasmid (Supplementary Figure 1). These regions were assessed for their percentage of GC skew using *genskew.csb.univie.ac.at* (Figure 4a). All were unwound rapidly and efficiently by FANCM:FAAP24 (Figure 4a). FANCM could also remove R-loops formed by transcription through the telomeric repeat sequence, otherwise known as TERRA transcripts (Figure 4b). These R-loops have previously been shown to have a strong G-quadraplex forming ability (Martadinata & Phan, 2013), suggesting that FANCM’s R-loop processing capability extends to R-loops that contain G-quadraplexes either within the displaced strand or between the RNA and the displaced strand (Arora *et al*, 2014), and supports a role for FANCM in the TERRA-dependent maintenance of telomeres by the ALT pathway of telomere maintenance (Lee *et al*, 2014, Pan *et al*, 2017).

**Figure 4:**
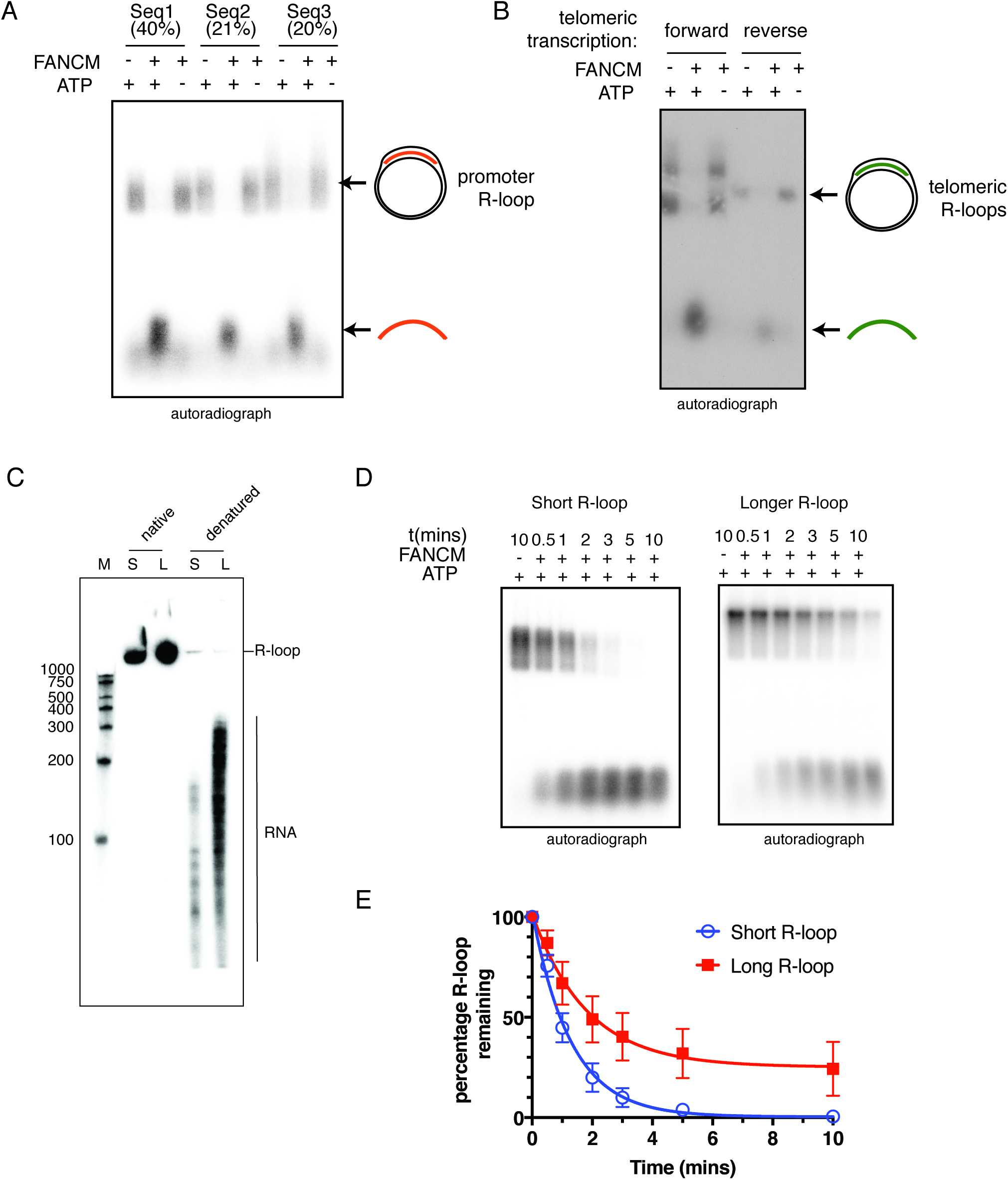
FANCM-FAAP24 can process R-loops irrespective of sequence, length or genomic origin. A) R-loop processing assay with three independently derived R-loop forming sequences. The GC skew % of each sequence is indicated in brackets above the autoradiograph. B) R-loops can be formed by *in vitro* transcription in a forward or reverse direction through a 1.4kb telomeric repeat sequence. FANCM:FAAP24 was tested against both products. C) R-loops of similar sequence but different length were generated (see materials and methods) and shown to produce trapped RNA species of different sizes on a PAGE gel. Short (S) and long (L) plasmid based R-loops are shown in native or denatured states. D-E) Substrates from C were incubated in a representative timecourse with FANCM:FAAP24 and R-loop remaining plotted from an average of 3 experiments (±stderr).

As R-loops of greater than 600bp can be detected in a cellular context (Ginno *et al*, 2012) we examined whether FANCM-FAAP24 could processively unwind R-loops containing different lengths of RNA. To do this we generated a plasmid containing ~1148bp of repeats of the immunoglobulin class switch sµ sequence (Supplementary Figure 1) downstream of a T7 promoter. We compared the size of RNA trapped within these R-loops to those found in the standard R-loop (consisting of a single sµ repeat of 143 bp) by subjecting the purified R-loops to Urea-PAGE. Denaturation revealed multiple RNA species of different lengths within the population of R-loops (Fig.4c). One explanation for this observation is that RNase A could cleave unpaired single strand regions within a longer R-loop, hence making the actual hybrid pieces appear shorter. Alternatively, T7 RNA polymerase could be stalling stochastically at different points along the template. Importantly though, the RNA fragments between the standard and long R-loops gave different average sizes of ~150 and ~300 nucleotides respectively (Figure 4c). We could therefore compare the rate of unwinding on R-loops of similar sequence but different length. Using the defined conditions of this assay FANCM-FAAP24 can unwind 100% of short plasmid R-loops within 5 minutes, with ~55% unwinding occurring within the first minute (Figure 4d). In contrast FANCM-FAAP24 takes 10 minutes to achieve ~85% unwinding of longer plasmid R-loops, a similar rate of unwinding when corrected for length (Figure 4d).

Collectively, these data further support the concept that FANCM:FAAP24 traverses long tracks of DNA without dissociation (i.e. processivity), to remove RNA molecules trapped within extended R-loop sequences.

### R-loop displacement is a conserved feature of FANCM family proteins

FANCM protein is a 230kDa protein, with an N-terminal translocase domain, a C-terminal ERCC4 structure specific DNA binding domain, and multiple protein:protein interaction domains that recruit additional DNA repair factors (Blackford *et al*, 2013, Coulthard *et al*, 2013, Deans & West, 2009) (Figure 5a). Only the N-terminal translocase domain is conserved in homologs from lower eukaryotes, such as Mph1 from *Saccharomyces cerevisiae*, and Fml1 from *S. pombe* (Whitby, 2010). RecG proteins are thought to be the closest relatives of FANCM/Mph1/Fml1 in bacteria (Gari *et al*, 2008a, Sun *et al*, 2008). To test whether R-loop unwinding is a conserved feature of these “FANCM family” members we purified recombinant *S.cerevesae* Mph1 *S.pombe* Fml1 or *Thermatoga maritima* RecG and tested their activity against our co-transcriptional R-loops (Figure 5b). RecG has previously been demonstrated to unwind synthetic linear R-loops (Vincent *et al*, 1996) but Mph1 and Fml1 have never been tested. Further data, collected using a FANCM fragment (aa 1-800), revealed the N-terminal conserved translocase domain of FANCM is sufficient for *in vitro* R-loop removal (Figure 5c). All of these enzymes displaced the co-transcriptional R-loops with similar activity, indicating a conserved function of FANCM-family proteins.

**Figure 5.**
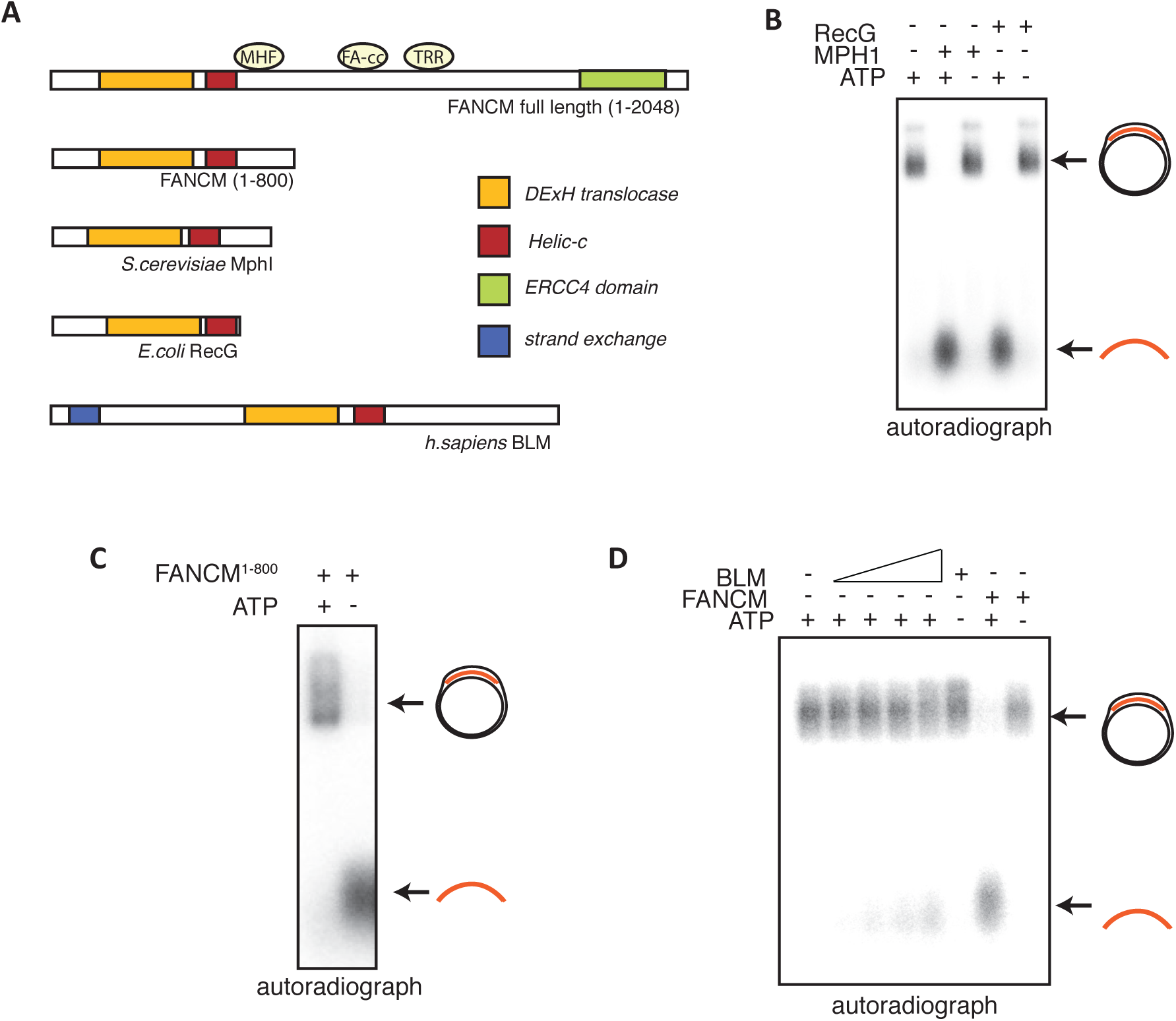
Conserved action of FANCM-like proteins in R-loop metabolism. A) Domain organization of FANCM orthologs, and BLM showing conserved domains. For FANCM, interaction sites are shown for: MHF=MHF1/2 complex, FAcc=Fanconi anemia core complex, TRR=Top3A-RMI1-RMI2 complex. B) MPH1 and RecG (1nM) unwind plasmid based R-loops in an ATP dependent manner, in a 10 minute reaction. C) FANCM (1nM) 1-800 retains the ability to unwind R-loops *in vitro*. D) Autoradiograph showing BLM unwinding plasmid based R-loops is ATP dependent but less efficient then FANCM. The concentrations of BLM used was 1, 10, 20, 40, 40nM (lane2-6) or 1nM for FANCM-FAAP24 (lane7-8).

FANCM protein (and Mph1 and RecG (McGlynn *et al*, 1997, Prakash *et al*, 2009)) can also unwind D-loops, which share structural similarities with R-loops (see Discussion). D-loop formation is an intermediate in DNA repair by homologous recombination which requires RAD51 or RecA recombinase. In addition to FANCM-family members, the RecQ helicase BLM (mutated in Bloom’s Syndrome) can unwind D-loops (Bachrati *et al*, 2006). We therefore tested BLM for its ability to unwind co-transcriptional R-loops. We carried out the assay with increasing amounts of BLM (Figure 5d). While BLM protein could unwind a very small fraction of the plasmid R-loops at high molar ratios of enzyme to plasmid (highest concentration 40nM, to 1nM substrate), it did this very slowly compared to FANCM:FAAP24, which rapidly unwound R-loops at stoichiometric and sub-stoichiometric concentrations (1nM for Figure 5d). In contrast to its much lower activity towards R-loop substrates, BLM was as efficient as FANCM in assays using a plasmid D-loop substrate (Supplemental Figure 4). These data suggest that BLM is capable of unwinding R-loop structures, but unlike for FANCM, they are not its preferred catalytic substrate.

### FANCM KO cells but not FANCL KO cells are sensitive to agents that induce R-loop stabilization

We recently demonstrated that FANCM knockout cells, or those expressing a translocation deficient FANCM^K117R^ mutant protein, accumulate excessive R-loops under normal cell culture growth conditions (Schwab *et al*, 2015). Several chemicals have also been shown to increase R-loop prevalence, each by a different mechanism. These include inhibitors of the spliceosome (that promote R-loops through retention of intronic sequences), topoisomerase 1 inhibitors such as topotecan (that promote R-loops by stalling transcription), and reactive aldehydes (unknown mechanism) (Powell *et al*, 2013, Schwab *et al*, 2015, Wan *et al*, 2015). We used isogenic FANCM KO, FANCL KO or parental HCT116 cells (previously characterized by Wang et al (Wang *et al*, 2013)) and tested their sensitivity to increasing concentrations of these R-loop promoting compounds. We found that FANCM deficient cells are particularly sensitive to topotecan and the spliceosome inhibitor pladienolide B, while FANCL-deficient cells are not (Figure 6a–b). Both FANCM and FANCL deficient cells are exquisitely sensitive to acetyl aldehyde, which also generates DNA interstrand crosslink damage in addition to R-loop accumulation.

**Figure 6.**
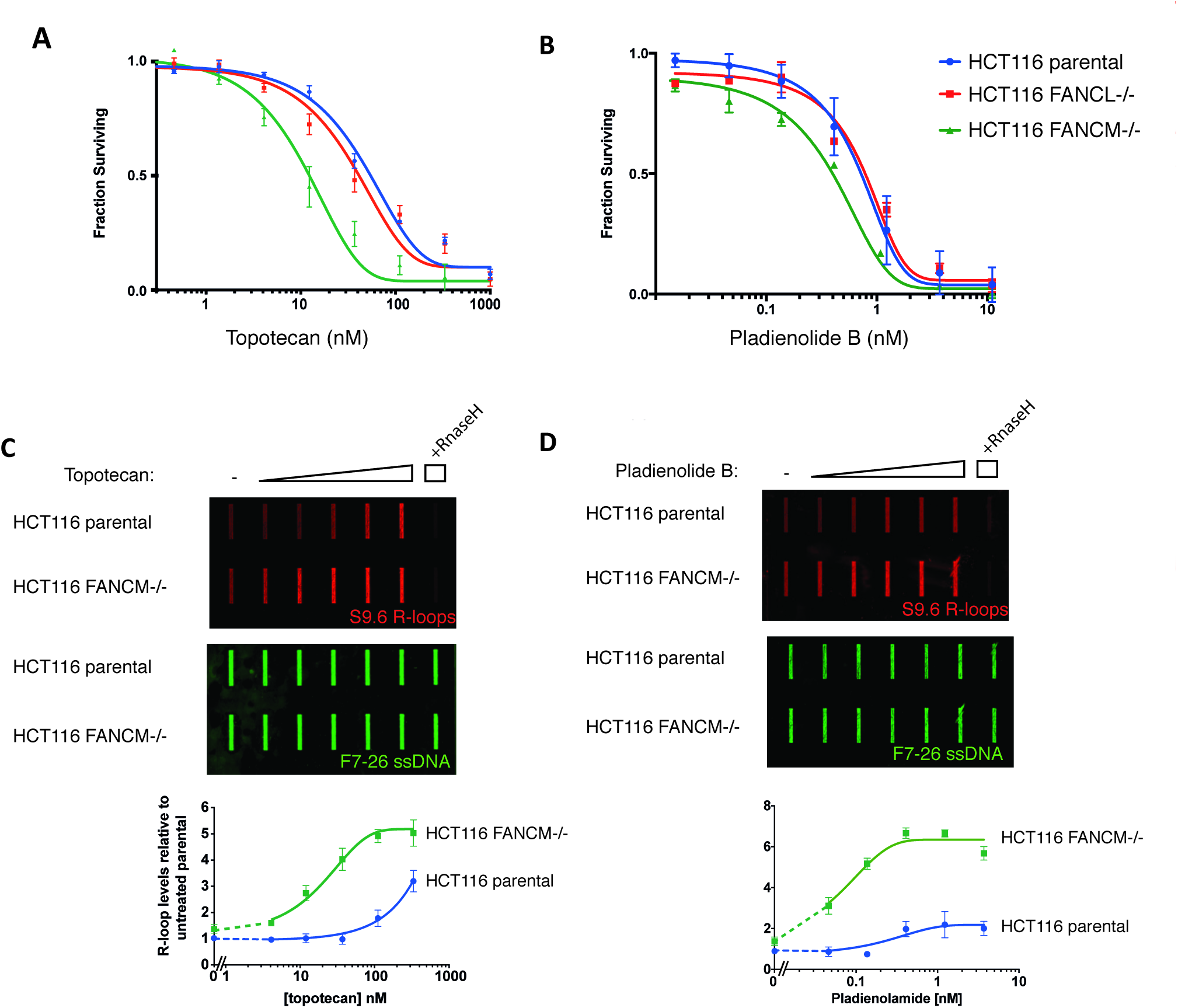
FANCM deficient cells are sensitive to R-loop stabilizing compounds. A-B) Dose response curves of parental and FANCM-/- HCT116 cell lines exposed to topotecan (left) or pladienolide B (right) for 72hours and stained with sulforhodamine B. C-D) Total R-loop levels measured by slot blotting of HCT116 genomic DNA extracted after increasing dose of topotecan or pladienolide B. Blots were probed with S9.6 anti-DNA:RNA hybrid or F7-26 anti-DNA, far-red secondary antibodies and detected by LiCor Odyssey imaging. RNaseH treated genomic DNA was used as a control for S9.6 DNA:RNA hybrid specificity. Quantification of S9.6 verses F7-26 slot blot signal from 3 experiments +/− sterr is shown in graphed form.

Importantly, sensitivity to both topotecan and pladienolide B correlated with an inflection point in total cellular R-loop levels in response to these drugs. This was measured by slot blot of genomic DNA probed with anti-DNA:RNA hybrid monoclonal antibody S9.6 (Figure 6c–d). R-loop levels and LD50 dosage were highly related, and suggests that cellular viability in response to these compounds correlates with a threshold R-loop level. This maximum tolerated level is reached with lower doses of either topotecan or pladienolide B in FANCM-deficient cells. This result indicates that R-loop metabolism requires FANCM activity independent of the other FA ICL repair proteins, to catalytically unwind R-loops formed under both physiological and drug-induced conditions.

## Discussion

Even though FANCM contains a SF2 helicase domain, exhaustive investigations have never uncovered a direct helicase activity of the protein against any DNA substrate (Coulthard *et al*, 2013, Gari *et al*, 2008a, Gari *et al*, 2008b, Meetei *et al*, 2005, Mosedale *et al*, 2005, Whitby, 2010, Xue et al, 2008). Instead, FANCM is thought to act on junctions in DNA as a branch point translocase (Gari *et al*, 2008a, Gari et al, 2008b). In this manner, it utilizes its ATP-dependent motor to reanneal DNA and further displace annealed strands ahead of the junction, without directly acting to unwind DNA like a helicase. This has been proposed for two different DNA structures: (i) stalled replication forks, whereby the annealing of nascent DNA strands by FANCM catalyzes replication fork reversal and the formation of a chicken foot structure and (ii) during recombination and D-loop formation, FANCM can catalyze displacement of the invading structure (Figure 7). Both of these activities have also been described for yeast Mph1 and bacterial RecG (McGlynn *et al*, 1997, Muller & West, 1994, Prakash *et al*, 2009). We propose a similar function for FANCM, Mph1 and RecG at R-loops at stalled transcription complexes. The branchpoint of all three structures, and the catalytic mechanism required for their translocation, is identical. In the case of stalled replication forks, FANCM creates the substrate necessary for resumption of replication (Gari *et al*, 2008a). But for D-loops and R-loops, FANCM is acting to suppress illegitimate recombination and/or barriers to DNA replication and transcription.

**Figure 7.**
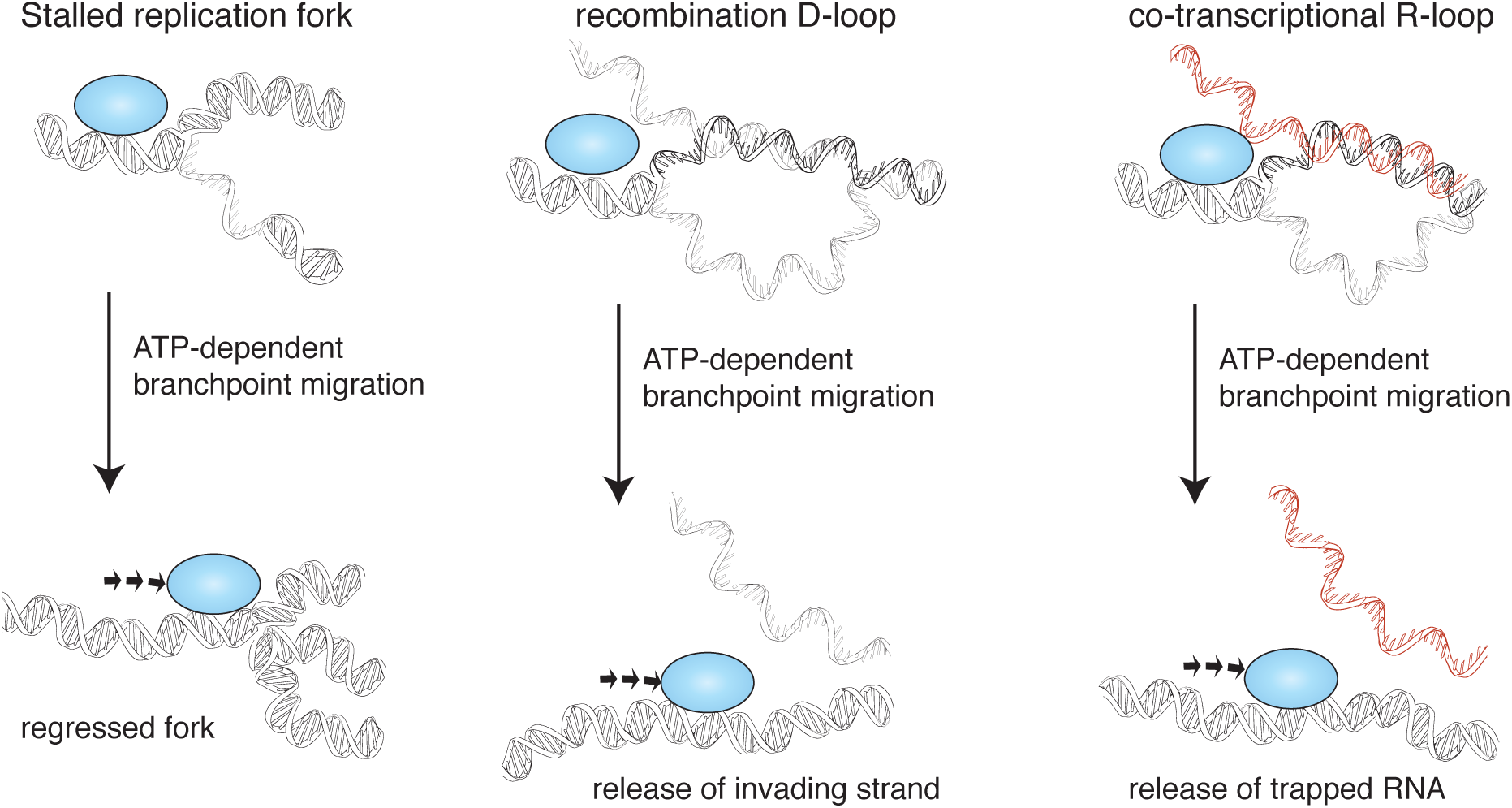
Model for FANCM-family-mediated maintenance of genome stability by a common mechanism of branchpoint translocation. Branchpoint translocation is indicated for 3 different DNA structures associated with genome instability. FANCM/Mph1/RecG or other family member (blue sphere in cartoon) binds specifically to junction structure in DNA. This engages the motor activity of the enzyme, to continuously push the junction. For a stalled replication fork (left) this leads to formation of a regressed fork, as the nascent DNA strands anneal. For a D-loop or R-loop, branchpoint migration leads to displacement of a ssDNA or RNA molecule respectively.

R-loop branch migration does not appear to be a general property of all translocases and helicases. For example, another SF2 helicase, BLM, can displace RNA from the R-loop, but only weakly when used at high concentration; similar to observations made for the bacterial enzyme RuvAB (Vincent *et al*, 1996). Like RuvAB, BLM is able to act to directly unwind DNA:RNA heteroduplexes constructed from synthetic oligonucleotides (Chang *et al*, 2017, Popuri *et al*, 2008), but these contained heterologous base-pairing, or extensive ssDNA regions that are not present in “native” co-transcriptional R-loops. The weak activity of BLM at high enzyme to substrate ratios (40:1, Figure 5d) on co-transcriptional R-loops, is probably the result of “accidental” helicase activity with the R-loop as the protein moves along double stranded DNA (Cheok *et al*, 2005), rather than the direct branch migration mechanism proposed for FANCM, Fml1, Mph1 and RecG. Future work should also compare the activity of FANCM to that of other proposed R-loop metabolizing enzymes such as Senataxin (Yuce & West, 2013), Aquarius (Sollier *et al*, 2014) and DHX9 (Chakraborty & Grosse, 2011). It should be noted that none of these enzymes in purified form have been directly tested for activity on co-transcriptional R-loops.

But what is the nature of R-loops that are acted upon by FANCM? R-loops can form at multiple different loci. Indeed, mathematical modeling suggests that every transcribed region is able to form an R-loop, but that increased observation at some loci comes from the fact that particular sequences are more prone to their formation, such as G-rich sequences (Belotserkovskii *et al*, 2017). Another consequence paradoxically, is that within a cell population, highly transcribed genes are more likely to be discovered in an R-loop-bound, arrested state. In this study, we have shown that FANCM can displace RNA from all co-transcriptional R-loops tested, including those that are long and those with very high G-content. FANCM therefore has the potential to act upon any R-loop in the genome. This might include R-loops required to initiate class switch recombination at the immunoglobulin heavy chain locus in activated B cells, which regulates immune function and occurs during G1 phase of the cell cycle (Schrader *et al*, 2007). Some FA mouse models show minor class switching defects (Nguyen *et al*, 2014), although *Fancm*-deficient mice have not been tested in this respect (Singh *et al*, 2009). We favor the hypothesis that FANCM’s R-loop regulation activity concerns their removal ahead of, or encountered by, the replication fork. This is because overwhelming evidence points to a role for the Fanconi anemia pathway in DNA damage during S phase (Deans & West, 2011) and the fact that FANCM protein is enriched in chromatin at sites of ongoing replication (Blackford *et al*, 2013, Castella *et al*, 2015). But DNA:RNA hybrids were also recently shown to form by transcription from dsDNA breaks during resection, when RNA polymerases become loaded onto broken DNA ends (Ohle *et al*, 2016). FANCM may play some role in displacing such hybrids in prevention of over-resection, another phenotype of FANCM-deficient cells (Blackford *et al*, 2013).

Finally, the maintenance of telomeres is also regulated by R-loops in some cancers (Azzalin *et al*, 2007). TERRA transcripts are produced by transcription of the C-rich telomeric DNA strand and are essential for the ALT mechanism of telomere maintenance by recombination. FANCM is necessary for ALT (Pan *et al*, 2017), and it is possible that this is because it acts on TERRA transcripts to permit telomere replication or somehow regulate telomere recombination. Our *in vitro* experiments demonstrate that FANCM displaces the G-quadraplex stabilized TERRA transcripts as efficiently as it acts on promoter- or switch-region- based R-loops.

While several physiological R-loops may be targeted by FANCM activity, it is also clear that chemically induced (or pathological) R-loops accumulate more rapidly, and for longer periods, in FANCM-deficient cells (Figure 6). In particular, R-loop promoting inhibitors of topoisomerase 1 or splicing promote R-loop accumulation at lower concentrations in FANCM-knockout cells, and this correlates with increased cell death induced by these drugs. Camptothecin sensitivity was also observed in cells from *Fancm*-deficient mice (Bakker *et al*, 2009), Mph1 deficient yeast (Scheller *et al*, 2000) and RecG deficient bacteria (Sutherland & Tse-Dinh, 2010). Such sensitivity is not a common property of homologous recombination deficiency, but is seen for a subset of HR proteins that also play a role in R-loop metabolism, such as BRCA1 and BRCA2 (Bhatia *et al*, 2014, Hill *et al*, 2014). Recent evidence suggests that transcription replication collisions promoted by splicing inhibitors or topoisomerase poisons could provide a major mechanism for the therapeutic action of these drugs (Sollier & Cimprich, 2015). As such, FANCM-deficiency or FANCM-overexpression could modulate the tumour and normal cellular response to these drugs in clinical use.

In conclusion, our biochemical reconstitution of co-transcriptional R-loop formation has established branch point translocation as a major mechanism of R-loop release. Given the emerging role of R-loops in cancer (Sollier & Cimprich, 2015, Stork *et al*, 2016), it is possible that a defect in FANCM mediated R-loop metabolism is directly responsible for the tumour prone phenotype of *FANCM*-associated Fanconi anaemia (homozygous mutations) and familial breast cancer (heterozygous carriers). The highly processive *in vitro* activity of FANCM, and strong R-loop phenotype after treatment with various chemotherapy drugs is consistent with an important role for processing of R-loops by FANCM in human disease.

## Methods

### Co-transcriptional R-loop plasmid design and construction

Co-transcriptional R-loop sequences for the human APOE or SNRPN promoter sequences or murine µ-switch repeat or AIRN promoter sequences were synthesized with a 5’-flanking T7 promotor and 3’-T7-terminator sequence and cloned into the EcoRI and HindIII sites of pUC19 (followed by sequence verification) by General Biosystems (Sequences are provided in Supplemental Figure 3). To generate the longer R-loop plasmid (pUC19 SR long), successive rounds of restriction cloning was undertaken to concatermerize the µ-switch repeat through repetitive subcloning of AvrII/HindIII digested fragments into SpeI/HindIII digested pUC19-SR. Plasmids were transformed into NEB 10-beta cells (NEB), plated and midi prepped (Qiagen). DNA concentration was established by nanodrop. R-loop forming plasmids have been deposited at Addgene. pcDNA6-Telo and TeloR contain a 0.8 kb fragment of human telomeric repeat cloned downstream of a T7 promoter (Arora *et al*, 2014).

### R-loop generation and purification

2µg of plasmid DNA was incubated in a final reaction volume of 200µl containing 1x T7 polymerase reaction buffer (NEB), 25 units of T7 polymerase (NEB), 2.25mM of each nucleotide CTP, GTP, ATP and 825nM UTP-ɑ-^32^P 3000 Ci/mmol (Perkin Elmer) for 1 hour at 37°C. The reaction was stopped by heat denaturation at 65°C for 20mins. 100µl of 1.05M NaCl and 0.03M MgCl_2_ buffer was added to each reaction plus 2.5µg of Rnase A (EpiCentre) for 1hr at 37°C. R-loops where then purified by 2x phenol/chloroform using phase lock tubes (Quanta Bio), precipitated in a final concentration of 0.3M Na Acetate and 70% ethanol at −20°C overnight. Next day the samples were centrifuged at 13,000 *x g* in table top centrifuge for 30min.

Supernatant was removed and samples were washed with 70% ethanol and centrifuged for a further 10min. Supernatant was removed and pellets were left to air dry. R-loops were resuspended in 10mM Tris pH8, then ran through 2x S-400 columns (GE Healthcare) to remove unincorporated nucleotides, quantified using nanodrop and stored at 4°C.

### Protein Purification

The following were purified as previously described: FLAG-FANCM-FAAP24 and FLAG-FANCM_K117R_-FAAP24 (Coulthard *et al*, 2013), RPA (Henricksen *et al*, 1994), RecG (Singleton *et al*, 2001) (a kind gift of Steve West, Francis Crick Institute), Flag-BLM, Flag-Mph1 and Flag-Topoisomerase IIIα-RMI1-RMI2 expression vectors were cloned into pFL or pUCDM baculovirus vectors and subsequently integrated into the Multibac Bacmid (Berger *et al*, 2004). For these proteins, 1L High 5 *Trichoplusia ni* cells (1 × 10^6^/ml, Invitrogen) were infected with virus (MOI=2.5). Cells were harvested 72 hours after infection at 500 x *g*, 4°C and pellets washed with 1xPBS. Cells were lysed on ice in 0.5M NaCl, 0.02M TEA pH 7.5, 1mM DTT, 10% glycerol plus mammalian protease inhibitors (Sigma P8340-5ml) and sonicated on ice 5x 10 sec bursts. Lysates was clarified by centrifugation at 35,000G for 40 minutes at 4°C. Clarified supernatant was then incubated with equilibrated Flag M2 resin (Sigma) for 1hr on a roller at 4°C. Flag resin was then subjected to 4x batch washes with lysis buffer (without mammalian protease inhibitors) with 5 minutes on a roller at 4°C between each spin. The resin was then placed into gravity flow column for a final wash and protein was eluted with 100µg/ml Flag peptide. Flag-Mph1 and Flag-FANCM-FAAP24 complexes were subjected to further purification by ssDNA affinity resin (Sigma): Flag elutions containing FANCM were pooled and diluted to have a final concentration of 100mM NaCl, 20mM TEA pH7.5, 10% glycerol, 1mM DTT (buffer B) and added to 400µl ssDNA resin overnight on a roller at 4°C. The resin was then placed down gravity flow column and washed with 10CV of buffer B. FANCM-FAAP24 complexes were eluted with buffer B containing 0.5M NaCl. Proteins were quantified using BSA titrations on SDS-PAGE gels. All proteins were flash frozen in their final buffers and stored at −80°C. Topoisomerase I from *E.coli* and all restriction enzymes were purchased from New England Biolabs.

### R-loop unwinding assays

R-loop unwinding reactions (10µl) contained 1nM of R-loop, 1mM ATP, 2µl of protein (protein concentrations stated in main text) in R-loop buffer (6.6mM Tris pH7.5, 3% glycerol, 0.1Mm EDTA, 1mM DTT, 0.5mM MgCl_2_) and incubated at 37°C for time as shown in figures. Reactions were stopped by adding 2µl of stop buffer (10mg.ml^-1^ proteinase K (NEB), 1% SDS) and incubated for 15min at 37°C. 2µl of 50% glycerol was added to samples prior to loading onto 1% or 0.8% agarose TAE gels, run at 100V in TAE buffer (40mM Tris, 20mM acetic acid, 1mM EDTA) for 60-90 mins. Gels were then crushed between precut biodyene B membranes (Pall) for 1hour, exposed overnight to a GE phosphor-screen and imaged on a Typhoo scanner (GE Biosciences). To visualize DNA, agarose gels were post stained with Sybr gold (Thermofisher) 1 in 10,000 in TAE.

Quantification of R-loop unwinding was performed using Image J and Prism software.

### Cell based assays

HCT116- or FANCM-/- and FANCL-/- derivatives were provided by Lei Li (University of Texas MD Anderson). Cell lines were authenticated by G-banding (St Vincent’s Cytogenetics) and maintained in DMEM + 10% fetal bovine serum at 37’C, 5% CO_2_ in a humidified chamber. For drug sensitivity assays, cells were plated in 96-well plates at 1,500 cells/well, then treated 24hrs later with various concentrations of topotecan (aka Hycamptin®, GSK) or pladienolamide B (Calbiochem). After 72hr, survival was measured using sulforhodamine B assay read at 550nm on a EnSpire Plate reader (Perkin Elmer).

To measure total cellular R-loop levels, HCT116 cells were treated with drug or vehicle for 4hr. Total genomic DNA was extracted using Isolate II kit (Bioline). 1µg of genomic DNA was slot blotted, using a BioRad Microfiltration apparatus, onto Biodyne B Nylon membrane (Thermo Fisher), which was then air-dried and blocked in Odyssey blocking buffer (LiCor). The membrane was then probed with 0.5µg/mL S9.6 anti-DNA:RNA monoclonal antibody (produced and purified in house from S9.6 hybridoma (ATCC)) and 10ng/ml anti-ssDNA (F7-26, Millipore). Atto800-anti-mouse (LiCor) and Cy5-conjugated anti-IgM antibody (Millipore) were used to visualize the level of DNA:RNA hybrids and total DNA detected by the primary antibodies, and visualized and quantified using Odyssey LiCor dual color imaging system and accompanying software.

## Author contributions

Conceptualization, C.H., J.J.O, and A.J.D.; Methodology, C.H., J.J.O., S.v.T., V.J.M., and A.J.D.; Investigation, all authors.; Writing - Original Draft, C.H. and A.J.D..; Writing - Review & Editing C.H., J.J.O, E.D. and A.J.D.; Funding Acquisition, A.J.D.; and Supervision, C.H., and A.J.D.

## Acknowledgements

We thank members of the Genome Stability Unit for input and discussions, and Jörg Heierhorst, Wayne Crismani and Wojciech Niedzwiedz for comments and suggestions on the manuscript. Thanks to Claus Azzalin for providing telomeric repeat plasmids, Lei Li for providing the HCT116 cell lines and Steve West for RecG. AJD is a Victorian Cancer Agency fellow, JJO received scholarship from the Leukaemia Foundation (Australia). This work was funded by National Health and Medical Research Council Australia (grants 1123100 and 1126004), Cancer Council of Victoria, Buxton trust and the Victorian Government’s OIS Program.

## References

Arora R, Lee Y, Wischnewski H, Brun CM, Schwarz T, Azzalin CM (2014) RNaseH1 regulates TERRA-telomeric DNA hybrids and telomere maintenance in ALT tumour cells. Nat Commun 5: 5220

Arudchandran A, Bernstein RM, Max EE (2004) Single-stranded DNA breaks adjacent to cytosines occur during Ig gene class switch recombination. J Immunol 173: 3223–9

Azzalin CM, Reichenbach P, Khoriauli L, Giulotto E, Lingner J (2007) Telomeric repeat containing RNA and RNA surveillance factors at mammalian chromosome ends. Science 318: 798–801

Bachrati CZ, Borts RH, Hickson ID (2006) Mobile D-loops are a preferred substrate for the Bloom's syndrome helicase. Nucleic Acids Research 34: 2269–2279

Bakker ST, van de Vrugt HJ, Rooimans MA, Oostra AB, Steltenpool J, Delzenne-Goette E, van der Wal A, van der Valk M, Joenje H, te Riele H, de Winter JP (2009) Fancm-deficient mice reveal unique features of Fanconi anemia complementation group M. Hum Mol Genet 18: 3484–95

Belotserkovskii BP, Soo Shin JH, Hanawalt PC (2017) Strong transcription blockage mediated by R-loop formation within a G-rich homopurine-homopyrimidine sequence localized in the vicinity of the promoter. Nucleic Acids Res

Berger I, Fitzgerald DJ, Richmond TJ (2004) Baculovirus expression system for heterologous multiprotein complexes. Nature biotechnology 22: 1583–7

Bhatia V, Barroso SI, Garcia-Rubio ML, Tumini E, Herrera-Moyano E, Aguilera A (2014) BRCA2 prevents R-loop accumulation and associates with TREX-2 mRNA export factor PCID2. Nature 511: 362–5

Blackford AN, Schwab RA, Nieminuszczy J, Deans AJ, West SC, Niedzwiedz W (2013) The DNA translocase activity of FANCM protects stalled replication forks. Hum Mol Genet 21: 2005–16

Castella M, Jacquemont C, Thompson EL, Yeo JE, Cheung RS, Huang JW, Sobeck A, Hendrickson EA, Taniguchi T (2015) FANCI Regulates Recruitment of the FA Core Complex at Sites of DNA Damage Independently of FANCD2. PLoS Genet 11: e1005563

Chakraborty P, Grosse F (2011) Human DHX9 helicase preferentially unwinds RNA-containing displacement loops (R-loops) and G-quadruplexes. DNA Repair (Amst) 10: 654–65

Chang EY, Novoa CA, Aristizabal MJ, Coulombe Y, Segovia R, Chaturvedi R, Shen Y, Keong C, Tam AS, Jones SJM, Masson JY, Kobor MS, Stirling PC (2017) RECQ-like helicases Sgs1 and BLM regulate R-loop-associated genome instability. J Cell Biol 216: 3991–4005

Cheok CF, Wu L, Garcia PL, Janscak P, Hickson ID (2005) The Bloom's syndrome helicase promotes the annealing of complementary single-stranded DNA. Nucleic Acids Res 33: 3932–41

Collis SJ, Ciccia A, Deans AJ, Horejsí Z, Martin JS, Maslen SL, Skehel JM, Elledge SJ, West SC, Boulton SJ (2008) FANCM and FAAP24 function in ATR-mediated checkpoint signaling independently of the Fanconi anemia core complex. Mol Cell 32: 313–24

Coulthard R, Deans A, Swuec P, Bowles M, Costa A, West S, Costa A, McDonald NQ (2013) Architecture and DNA recognition elements of the Fanconi anemia FANCM-FAAP24 complex. Structure 21: 1648–1658

Crismani W, Girard C, Froger N, Pradillo M, Santos JL, Chelysheva L, Copenhaver GP, Horlow C, Mercier R (2012) FANCM limits meiotic crossovers. Science 336: 1588–90

Deans AJ, West SC (2009) FANCM connects the genome instability disorders Bloom's Syndrome and Fanconi Anemia. Mol Cell 36: 943–53

Deans AJ, West SC (2011) DNA interstrand crosslink repair and cancer. Nat Rev Cancer 11: 467–80

El Hage A, French SL, Beyer AL, Tollervey D (2010) Loss of Topoisomerase I leads to R-loop-mediated transcriptional blocks during ribosomal RNA synthesis. Genes Dev 24: 1546–58

Garcia-Rubio ML, Perez-Calero C, Barroso SI, Tumini E, Herrera-Moyano E, Rosado IV, Aguilera A (2015) The Fanconi Anemia Pathway Protects Genome Integrity from R-loops. PLoS Genet 11:e1005674

Gari K, Décaillet C, Delannoy M, Wu L, Constantinou A (2008a) Remodeling of DNA replication structures by the branch point translocase FANCM. Proc Natl Acad Sci USA

Gari K, Decaillet C, Stasiak AZ, Stasiak A, Constantinou A (2008b) The Fanconi anemia protein FANCM can promote branch migration of Holliday junctions and replication forks. Mol Cell 29: 141–8

Ginno PA, Lim YW, Lott PL, Korf I, Chedin F (2013) GC skew at the 5' and 3' ends of human genes links R-loop formation to epigenetic regulation and transcription termination. Genome Res 23: 1590–600

Ginno PA, Lott PL, Christensen HC, Korf I, Chedin F (2012) R-loop formation is a distinctive characteristic of unmethylated human CpG island promoters. Mol Cell 45: 814–25

Hamperl S, Bocek MJ, Saldivar JC, Swigut T, Cimprich KA (2017) Transcription-Replication Conflict Orientation Modulates R-Loop Levels and Activates Distinct DNA Damage Responses. Cell 170: 774–786e19

Helmrich A, Ballarino M, Tora L (2011) Collisions between replication and transcription complexes cause common fragile site instability at the longest human genes. Mol Cell 44: 966–77

Henricksen LA, Umbricht CB, Wold MS (1994) Recombinant replication protein A: expression, complex formation, and functional characterization. J Biol Chem 269: 11121–32

Hill SJ, Rolland T, Adelmant G, Xia X, Owen MS, Dricot A, Zack TI, Sahni N, Jacob Y, Hao T, McKinney KM, Clark AP, Reyon D, Tsai SQ, Joung JK, Beroukhim R, Marto JA, Vidal M, Gaudet S, Hill DEet al. (2014) Systematic screening reveals a role for BRCA1 in the response to transcription-associated DNA damage. Genes Dev 28: 1957–75

Hong X, Cadwell GW, Kogoma T (1995) Escherichia coli RecG and RecA proteins in R-loop formation. EMBO J 14: 2385–92

Huertas P, Aguilera A (2003) Cotranscriptionally formed DNA:RNA hybrids mediate transcription elongation impairment and transcription-associated recombination. Mol Cell 12: 711–21

Lee M, Hills M, Conomos D, Stutz MD, Dagg RA, Lau LM, Reddel RR, Pickett HA (2014) Telomere extension by telomerase and ALT generates variant repeats by mechanistically distinct processes. Nucleic Acids Res 42: 1733–46

Madireddy A, Kosiyatrakul ST, Boisvert RA, Herrera-Moyano E, Garcia-Rubio ML, Gerhardt J, Vuono EA, Owen N, Yan Z, Olson S, Aguilera A, Howlett NG, Schildkraut CL (2016) FANCD2 Facilitates Replication through Common Fragile Sites. Mol Cell 64: 388–404

Martadinata H, Phan AT (2013) Structure of human telomeric RNA (TERRA): stacking of two G- quadruplex blocks in K(+) solution. Biochemistry 52: 2176–83

McGlynn P, Al-Deib AA, Liu J, Marians KJ, Lloyd RG (1997) The DNA replication protein PriA and the recombination protein RecG bind D-loops. J Mol Biol 270: 212–21

Meetei AR, Medhurst AL, Ling C, Xue Y, Singh TR, Bier P, Steltenpool J, Stone S, Dokal I, Mathew CG, Hoatlin M, Joenje H, de Winter JP, Wang W (2005) A human ortholog of archaeal DNA repair protein Hef is defective in Fanconi anemia complementation group M. Nat Genet 37:958–63

Mosedale G, Niedzwiedz W, Alpi A, Perrina F, Pereira-Leal JB, Johnson M, Langevin F, Pace P, Patel KJ (2005) The vertebrate Hef ortholog is a component of the Fanconi anemia tumor-suppressor pathway. Nat Struct Mol Biol 12: 763–71

Muller B, West SC (1994) Processing of Holliday junctions by the Escherichia coli RuvA, RuvB, RuvC and RecG proteins. Experientia 50: 216–22

Nakamura H, Oda Y, Iwai S, Inoue H, Ohtsuka E, Kanaya S, Kimura S, Katsuda C, Katayanagi K, Morikawa K, et al. (1991) How does RNase H recognize a DNA.RNA hybrid? Proc Natl Acad Sci U S A 88: 11535–9

Nguyen HD, Yadav T, Giri S, Saez B, Graubert TA, Zou L (2017) Functions of Replication Protein A as a Sensor of R Loops and a Regulator of RNaseH1. Mol Cell 65: 832–847.e4

Nguyen TV, Riou L, Aoufouchi S, Rosselli F (2014) Fanca deficiency reduces A/T transitions in somatic hypermutation and alters class switch recombination junctions in mouse B cells. J Exp Med 211: 1011–8

Ohle C, Tesorero R, Schermann G, Dobrev N, Sinning I, Fischer T (2016) Transient RNA-DNA Hybrids Are Required for Efficient Double-Strand Break Repair. Cell 167: 1001–1013.e7

Pan X, Drosopoulos WC, Sethi L, Madireddy A, Schildkraut CL, Zhang D (2017) FANCM, BRCA1, and BLM cooperatively resolve the replication stress at the ALT telomeres. Proc Natl Acad Sci U S A

Popuri V, Bachrati CZ, Muzzolini L, Mosedale G, Costantini S, Giacomini E, Hickson ID, Vindigni A (2008) The Human RecQ helicases, BLM and RECQ1, display distinct DNA substrate specificities. Biol Chem 283: 17766–76

Powell WT, Coulson RL, Gonzales ML, Crary FK, Wong SS, Adams S, Ach RA, Tsang P, Yamada NA, Yasui DH, Chedin F, LaSalle JM (2013) R-loop formation at Snord116 mediates topotecan inhibition of Ube3a-antisense and allele-specific chromatin decondensation. Proc Natl Acad Sci U S A 110: 13938–43

Prakash R, Satory D, Dray E, Papusha A, Scheller J, Kramer W, Krejci L, Klein H, Haber JE, Sung P, Ira G (2009) Yeast Mph1 helicase dissociates Rad51-made D-loops: implications for crossover control in mitotic recombination. Genes Dev 23: 67–79

Roy D, Yu K, Lieber MR (2008) Mechanism of R-Loop Formation at Immunoglobulin Class Switch Sequences. Molecular and Cellular Biology 28: 50–60

Scheller J, Schurer A, Rudolph C, Hettwer S, Kramer W (2000) MPH1, a yeast gene encoding a DEAH protein, plays a role in protection of the genome from spontaneous and chemically induced damage. Genetics 155: 1069–81

Schrader CE, Guikema JE, Linehan EK, Selsing E, Stavnezer J (2007) Activation-induced cytidine deaminase-dependent DNA breaks in class switch recombination occur during G1 phase of the cell cycle and depend upon mismatch repair. J Immunol 179: 6064–71

Schwab RA, Nieminuszczy J, Shah F, Langton J, Lopez Martinez D, Liang CC, Cohn MA, Gibbons RJ, Deans AJ, Niedzwiedz W (2015) The Fanconi Anemia Pathway Maintains Genome Stability by Coordinating Replication and Transcription. Mol Cell 60: 351–61

Shin JH, Kelman Z (2006) The replicative helicases of bacteria, archaea, and eukarya can unwind RNA-DNA hybrid substrates. J Biol Chem 281: 26914–21

Singh TR, Bakker ST, Agarwal S, Jansen M, Grassman E, Godthelp BC, Ali AM, Du CH, Rooimans MA, Fan Q, Wahengbam K, Steltenpool J, Andreassen PR, Williams DA, Joenje H, de Winter JP, Meetei AR (2009) Impaired FANCD2 monoubiquitination and hypersensitivity to camptothecin uniquely characterize Fanconi anemia complementation group M. Blood 114: 174–80

Singleton MR, Scaife S, Raven ND, Wigley DB (2001) Crystallization and preliminary X-ray analysis of RecG, a replication-fork reversal helicase from Thermotoga maritima complexed with a three-way DNA junction. Acta crystallographica Section D, Biological crystallography 57: 1695–6

Sollier J, Cimprich KA (2015) Breaking bad: R-loops and genome integrity. Trends Cell Biol 25:514–22

Sollier J, Stork CT, Garcia-Rubio ML, Paulsen RD, Aguilera A, Cimprich KA (2014) Transcription-coupled nucleotide excision repair factors promote R-loop-induced genome instability. Mol Cell 56: 777–85

Stork CT, Bocek M, Crossley MP, Sollier J, Sanz LA, Chedin F, Swigut T, Cimprich KA (2016) Co-transcriptional R-loops are the main cause of estrogen-induced DNA damage. eLife 5

Sun W, Nandi S, Osman F, Ahn JS, Jakovleska J, Lorenz A, Whitby MC (2008) The FANCM ortholog Fml1 promotes recombination at stalled replication forks and limits crossing over during DNA double-strand break repair. Mol Cell 32: 118–28

Sutherland JH, Tse-Dinh YC (2010) Analysis of RuvABC and RecG involvement in the escherichia coli response to the covalent topoisomerase-DNA complex. J Bacteriol 192: 4445–51

Szczelkun MD, Tikhomirova MS, Sinkunas T, Gasiunas G, Karvelis T, Pschera P, Siksnys V, Seidel R (2014) Direct observation of R-loop formation by single RNA-guided Cas9 and Cascade effector complexes. Proc Natl Acad Sci U S A 111: 9798–803

Vincent SD, Mahdi AA, Lloyd RG (1996) The RecG branch migration protein of Escherichia coli dissociates R-loops. J Mol Biol 264: 713–21

Wan Y, Zheng X, Chen H, Guo Y, Jiang H, He X, Zhu X, Zheng Y (2015) Splicing function of mitotic regulators links R-loop-mediated DNA damage to tumor cell killing. J Cell Biol 209: 235–46

Wang Y, Leung Justin W, Jiang Y, Lowery Megan G, Do H, Vasquez Karen M, Chen J, Wang W, Li L (2013) FANCM and FAAP24 Maintain Genome Stability via Cooperative as Well as Unique Functions. Molecular cell 49: 997–1009

Whitby MC (2010) The FANCM family of DNA helicases/translocases. DNA Repair (Amst) 9: 224–36

Wilson-Sali T, Hsieh T-s (2002) Preferential cleavage of plasmid-based R-loops and D-loops by Drosophila topoisomerase IIIβ. Proceedings of the National Academy of Sciences 99: 7974–7979

Xue Y, Li Y, Guo R, Ling C, Wang W (2008) FANCM of the Fanconi anemia core complex is required for both monoubiquitination and DNA repair. Hum Mol Genet

Yuce O, West SC (2013) Senataxin, defective in the neurodegenerative disorder ataxia with oculomotor apraxia 2, lies at the interface of transcription and the DNA damage response. Mol Cell Biol 33: 406–17

